# Mechanosensing by Piezo1 regulates osteoclast differentiation via PP2A-Akt axis in periodontitis

**DOI:** 10.1101/2024.09.04.611049

**Authors:** Satoru Shindo, Shin Nakamura, Mohamad Rawas-Qalaji, Alireza Heidari, Maria Rita Pastore, Motoki Okamoto, Maiko Suzuki, Manuel Salinas, Dmitriy Minond, Alexander Bontempo, Mark Cayabyab, Yingzi Yang, Janet L Crane, Maria Hernandez, Saynur Vardar, Patrick Hardigan, Xiaozhe Han, Steven Kaltman, Toshihisa Kawai

## Abstract

Mechanical stimulus to the multicellular bone unit (MBU) plays a key role in normal bone remodeling, whereas disuse osteoporosis, for example, represents loss of bone owing to lack of mechanical stresses. The analogy can be applied to a variety of pathogenic bone lytic complications, including periodontitis, in which local mechanical stress appears to be diminished. The activation of mechanosensitive Piezo1 Ca^2+^ channel expressed by osteoblasts and osteocytes in the MBU elicits the osteogenic signals in those cells. However, since osteoclast (OC)-specific Piezo1-gene knockout mice showed no skeletal phenotype, it has been assumed that Piezo1 might not play any role in OC-mediated bone remodeling. Here, however, we showed that mechanical stimulation of Piezo1 expressed on preosteoclasts (pre-OCs) downmodulates OC formation and, hence, bone resorptive activity in periodontitis, accompanied by significantly reduced expression of NFATc1, a master transcription factor for RANKL-induced OC-genesis. We know that the Ca^2+^/calcineurin/NFAT axis upregulates NFATc1 activation in pre-OCs. Interestingly, Piezo1-elicited Ca^2+^ influx did not affect NFATc1 expression. Instead, PP2A-mediated dephosphorylation of Akt downregulated NFATc1 in Piezo1-activated pre-OCs. However, systemic administration with Yoda1, a Piezo1 chemical agonist, or local injection of PP2A agonist, significantly downregulated the bone resorption induced in a mouse model of periodontitis, together with reduced numbers of TRAP^+^/phospho-Akt^+^ pre-OCs in local bone. These results suggest that mechanosensing by Piezo1 expressed on pre-OCs can downmodulate the RANKL-induced OC-genesis via the PP2A/Akt-dephosphorylation pathway, but that such Piezo1-mediated downregulation of bone resorption is attenuated in periodontitis.

**Significance Statement:** The mechanosensitive Ca^2+^ channel Piezo1 plays important regulatory roles in a variety of cellular activities. RANKL-mediated OC-genesis requires permissive co-stimulatory signal from ITAM receptors, such as OSCAR and TREM2, to trigger the calcineurin/calmodulin signaling axis via Ca^2+^ oscillation, thereby upregulating NFATc1 expression. Activation of Piezo1 remarkably suppressed RANKL-induced NFATc1 activation which, in turn, reduced OC-genesis. Such mechanical activation of Piezo1 expressed on pre-OCs induced intracellular Ca^2+^ influx. Nonetheless, PP2A-mediated dephosphorylation of Akt, not the calcineurin/calmodulin pathway, suppressed NFATc1 in RANKL-elicited OC-genesis and resultant bone resorption, both *in vitro* and *in vivo*. These results indicate that mechanostress applied to pre-OCs can downregulate pathogenic OC-genesis and that Piezo1, as the mediator, is a novel molecular target for the development of anti-osteolytic therapies.

## Introduction

Onset and progression of periodontitis result from overactivation of host immune response against opportunistic pathogens in the periodontal microbiome. This, in turn, leads to tissue-destructive inflammation of the periodontium, including alveolar bone resorption (1–3). Inflammation in periodontium causes angiogenesis, vasodilation and vascular permeability, allowing for the expanded migration of immune cells, including monocytes and macrophages, following T lymphocytes (4). Osteoclasts (OCs) act as the key player of bone resorption in inflammatory bone lytic diseases, such as rheumatoid arthritis (5, 6) and periodontitis (7, 8). Preosteoclasts (pre-OCs) are tartrate-resistant acid phosphatase (TRAP)^+^ mononucleated pre-OCs which fuse to form multinucleated mature TRAP^+^ OCs (9) by activation of the master transcription factor (TF) Nuclear factor of activated T-cells, cytoplasmic 1 (NFATc1) in an Receptor activator of nuclear factor kappa-β ligand (RANKL)-dependent manner (10). RANKL-mediated OC-genesis requires permissive costimulatory signaling from immunoreceptor tyrosine-based activation motif (ITAM) receptors (signaling transducers), such as Osteoclast-associated receptor (OSCAR) and Triggering receptor expressed on myeloid cells 2 (TREM2), to trigger the calcineurin/calmodulin signaling axis via Ca^2+^ oscillation (11, 12), in turn upregulating NFATc1 expression.

Monocytes in the circulation, a significant source of pre-OCs (13–16), adhere to capillary endothelium and migrate into the homeostatic bone remodeling site as well as bone lytic lesion upon receiving signals from lipid mediators and specific chemokines, including Sphingosine 1-phosphate, monocyte chemoattractant protein-1 (MCP-1), C-X-C motif chemokine 12 (CXCL12), stromal cell-derived factor-1 (SDF-1), C-X3-C motif ligand 1 (CX3CL1) or macrophage migration inhibitory factor (MIF) (17–22). Blood flow acts as a mechanical force on blood cells, such as T lymphocytes and platelets, as well as pre-OCs (23–26). Even though chronic inflammation in periodontitis causes expansion of capillary diameter (27), blood velocity in the vasculature of periodontitis lesion is significantly diminished (28). It is also reported that the rate of blood flow in healthy gingival tissue is significantly lower in the elderly than that of younger groups (29). To avoid the induction of pain at the periodontally diseased tooth, mastication at the affected tooth is also reduced (30, 31). Based on these lines of evidence, It can be posited that hematopoietic cells, including pre-osteoclasts (pre-OCs), in the periodontium affected by periodontitis are subjected to reduced mechanical stress compared to those in healthy periodontal tissue. Increased mechanical stimuli can alter various activities of cells in periodontium, including periodontal ligament cells and osteoblasts, (32, 33); outcomes of the opposite condition whereby the periodontium receives less mechanical stress, especially alveolar bone, are largely unknown in the context of periodontitis. Several studies have reported the suppressive effect of shear stress on *in vitro* OC-genesis, using mouse pre-OC RAW264.7 cells (34, 35). Nonetheless, it is unclear if mechanical stress also affects the OC-genesis of primary pre-OCs, both *in vitro* and *in vivo*. Furthermore, the molecular mechanism underlying such suppressive effect of mechanical stress on OC-genesis remains to be elucidated.

Therefore, to shed more light on these unresolved questions, we turned to the mechanosensory system, the importance of which was highlighted by the 2021 Nobel Prize awarded for the discovery of mechanosensitive Piezo Ca^2+^ channels (36). Specifically, upon mechanical stimulation, the Piezo1 channel is opened by rearrangement of cytoskeletal actin anchored to the proximal of Piezo1, thereby eliciting Ca^2+^ influx which induces cell signaling for a variety of activities (37–42). Here, we report that Piezo1 is the major mechanosensory receptor expressed on pre-OCs and that diminished mechanostress to pre-OCs can impede the Piezo1-mediated downmodulation of OC-genesis in periodontitis. We further discovered that activation of Piezo1 expressed on pre-OCs elicits a unique cell signal axis involving PP2A-mediated dephosphorylation of Akt which, in turn, suppresses the expression of NFATc1, a master TF for RANKL-induced OC-genesis.

## Results

### 1. Blood flow was reduced in the mouse periodontal with periodontitis

Although, as noted above, blood flow rate in the vasculature of human periodontitis lesion is significantly diminished (28), it is unknown whether blood flow in the periodontitis induced in mice is also reduced. To induce periodontitis, a silk ligature was attached to the second maxillary molar for 7 days which resulted in induction of local inflammation, represented by the elevated expression of TNF-α and IL-1β mRNAs (Fig. 1A). To assess the impact of periodontitis on the local blood flow, the blood perfusion unit (BPU) of palatal mesial or distal gingival tissue was measured using Laser Doppler Flowmetry. Irrespective of inflammation induced in the periodontal tissue, the BPU in periodontally diseased tissue was significantly reduced compared to that in healthy tissue (Fig. 1B and C).

**Figure 1.**
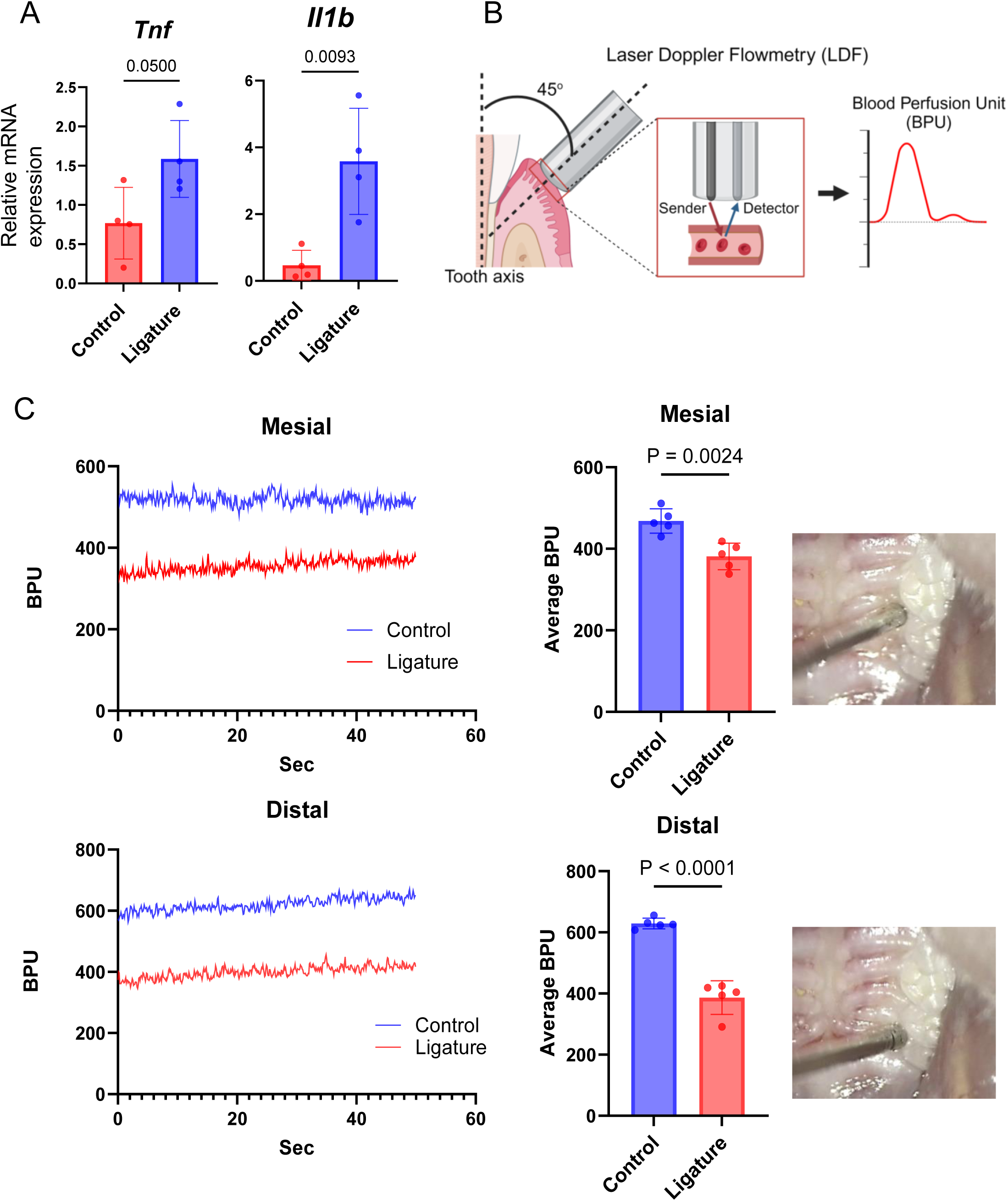
Induction of periodontitis in mouse reduced local BPU. (A) 5-0 silk were placed at upper second molar to induce murine periodontitis. The gene expression of inflammatory cytokines, including *Tnf* and *Il1b*, in gingival tissue from either periodontally affected or healthy gingiva was monitored by qPCR. (B) Real time BPU that reflects the flow rate of local vasculature in the palatal gingival tissue of live mouse was measured by laser doppler flowmetry as shown. BPU was measured at the mesial and distal sites of second maxillary molar to which a silk ligature was attached for induction of periodontitis. The inserted pictures show the control second maxillary molar that was not attached with ligature. Continuous detections of temporal change of real time BPU (50 s) were displayed. (C) Results in the bar graph were presented as the means ± SD.

### 2. Mechanical stress suppressed RANKL-induced OC-genesis *in vitro*

The Fluid oscillation caused by a microfluidics system was employed to evaluate whether RANKL-induced OC-genesis is influenced by mechanical stress. According to the previous report, blood flow rate in the microcapillaries of heathy periodontal tissue is approximately 20 dyn/cm^2^, but that in diseased periodontal tissue is diminished to approximately 5 dyn/cm^2^ (28). OC-genesis as well as OC-related gene expression (*Ocstamp*, *Mmp9*, *Acp5*, *Oscar*, *Dcstamp* and *Nfatc1*) were significantly suppressed by high flow rate (20 dyn/cm^2^) compared to low flow rate (5 dyn/cm^2^) or static condition (Fig. 2B and C). Moreover, shear flow generated in the tissue culture plate by a rocker (15°, 30 rpm) also suppressed RANKL-induced OC-genesis (Fig. S1A).

**Figure 2.**
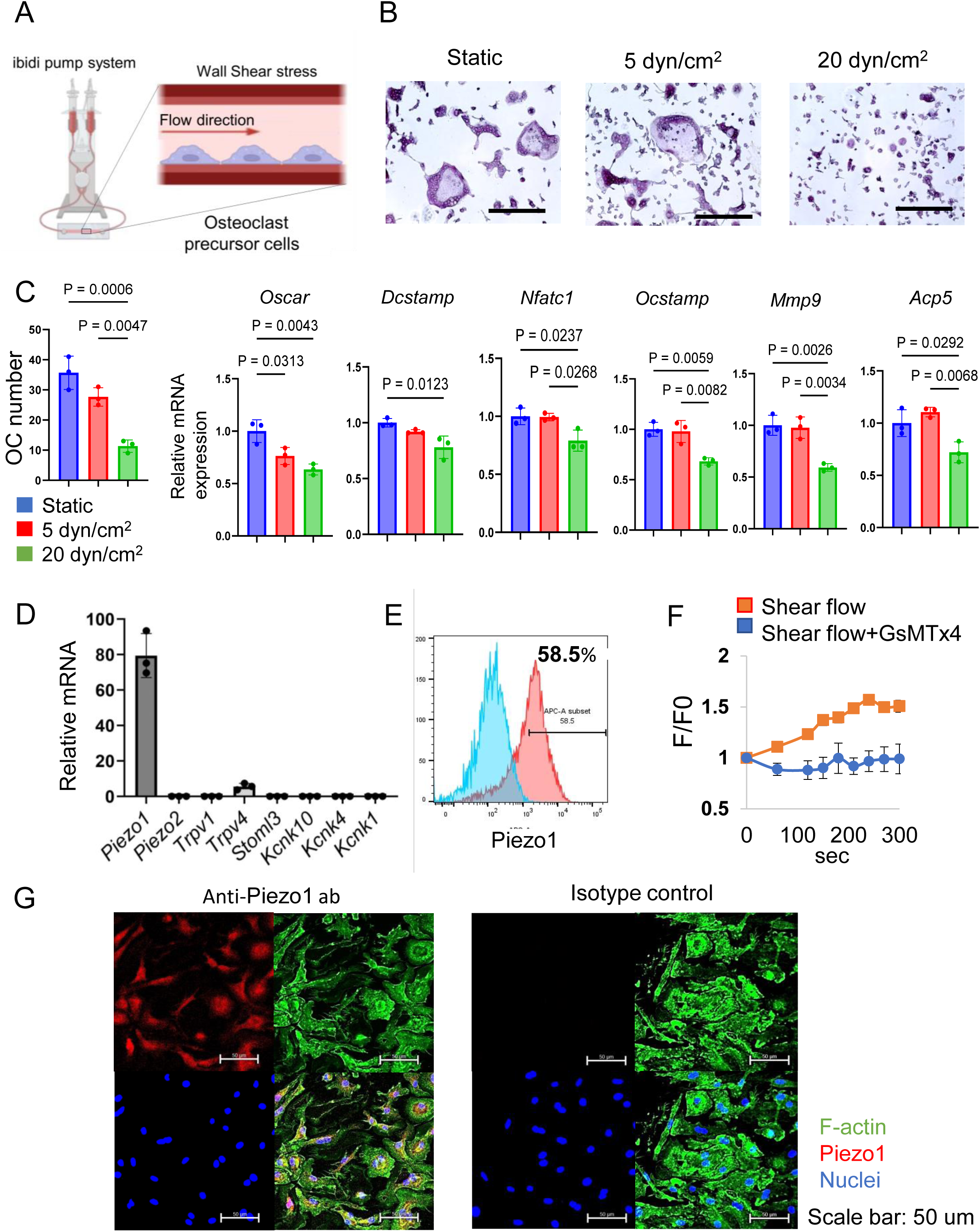
Shear stress inhibited OC-genesis, and pre-OCs expressed Piezo1 prominently. (A) Schematic image of stimulation by shear stress on OCs using the microfluidics system. (B and C) MCSF (25 ng/ml)-primed murine bone marrow-derived mononuclear cells were employed as pre-OCs. RANKL (10 ng/ml)-mediated OC-genesis with or without shear stress by microfluidics system was evaluated by TRAP staining and qPCR to monitor *Oscar*, *Dcstamp*, *Nfatc1 Ocstamp*, *Mmp9* and *Acp5*. (D) Expression of mechanosensory receptors, including *Piezo1*, *Piezo2, Trpv1, Trpv4, Stmol3, Kcnk10, Kcnk1* and *Kcnk4*, in pre-OCs was determined by qPCR. (E and G) Piezo1 protein expression was detected by immunofluorescence and flow cytometry, respectively. (F) Fluo-8-treated pre-OCs with or without GsMTx4 (1 uM) were stimulated by shear flow at 20 dyn/cm^2^ using the Bioflux microfluidics system. Ca^2+^ influx was analyzed with the Bioflux system. Data represent the mean ± SD of three independent experiments.

### 3. Piezo1 expressed on pre-OC functioned as a mechanosensory Ca^2+^ channel

Out of 8 major mechanoreceptors (*Piezo1*, *Piezo2*, *Trpv1*, *Trpv4*, *Stoml3*, *Kcnk10*, *Kcnk4* and *Kcnk1*), Piezo1 mRNA was the highest expressed. (Fig. 2D). Protein expression of Piezo1 in pre-OCs was confirmed by both immunofluorescence staining and flow cytometry (58.5 %) (Fig. E and G). GsMTx4, a spider venom that selectively inhibits Piezo1 (43–45), suppressed Ca^2+^ influx induced by microfluidics-generated shear flow in pre-OCs (Fig. 2G), indicating that pre-OCs appeared to sense shear flow-generated mechanical force via Piezo1.

### 4. Pharmacological Piezo1 activator inhibited OC-genesis and function

Yoda1 is a chemical agonist that can selectively open Piezo1 and promote intercellular Ca^2+^ to initiate a variety of biological events (46–49). We therefore used Yoda1 to determine if RANKL-induced OC-genesis was regulated by Piezo1. We found that TRAP-positive multinucleated OC formation, as well as bone resorptive activity, were both significantly diminished by Yoda1 administration (Fig. 3A). In addition, GsMTx4 inhibited Yoda1-induced Ca^2+^ influx in pre-OCs (Fig. S1B). To test the effect of Yoda1 on human OC-genesis, peripheral blood mononuclear cells (PBMC)-derived pre-OCs were employed. Yoda1-mediated Piezo1 activation also inhibited RANKL-mediated human OC-genesis (Fig. S2).

**Figure 3.**
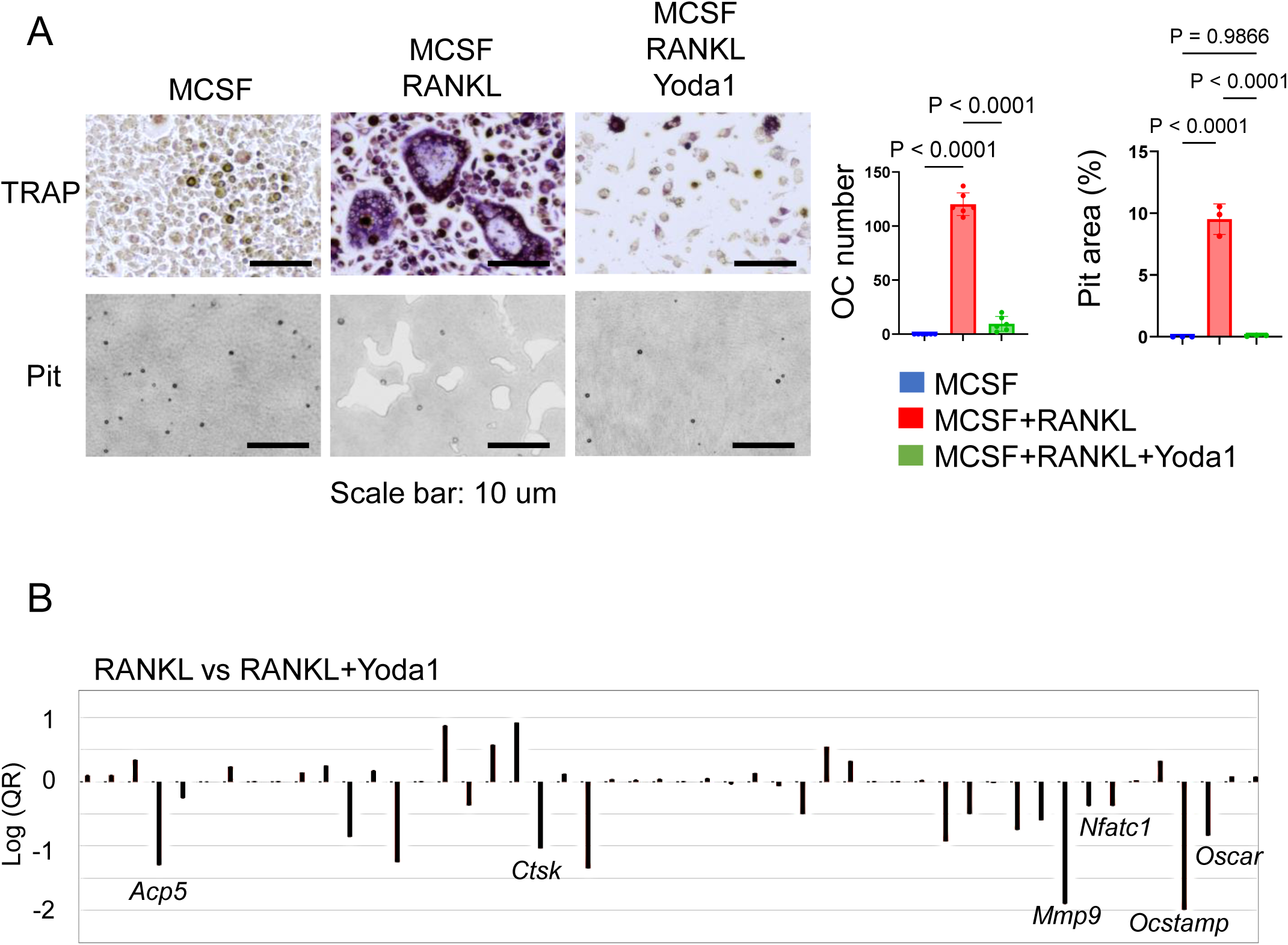
Pharmacological Piezo1 activator inhibited OC-genesis and function. (A) Pre-OCs were treated with RANKL (10 ng/ml) in the presence or absence of Yoda1 (5 µM) or vehicle control. After 6days, TRAP-positive OCs with three or more nuclei were counted as mature OCs. Pit formation activity by OCs was evaluated with imaging and calculation using Image J. (B) PCR array was performed to identify osteoclast-related genes regulated by Yoda1. Data represent the mean ± SD of three independent experiments.

PCR array was used to screen for RANKL-stimulated genes that were impaired by Yoda (Fig. 3B). Yoda1-mediated suppression of genes (*Ocstamp*, *Ctsk*, *Mmp9*, *Acp5* and *Oscar*) was also confirmed by qPCR (Fig. 4A). Upon stimulation with RANKL, NFATc1, a master TF of OC-genesis (10, 50, 51), translocates from cytoplasm to nucleus and induces the transcription of genes required for OC-genesis and fusion (52, 53). Yoda1 inhibited the expression of *Nfatc1* gene and its protein during RANKL-induced OC-genesis (Fig. 4B, C, D and Fig. S6). These results suggested that the pharmacological activation of Piezo1 caused the downregulation of RANKL-induced OC-genesis in conjunction with the suppression of both *Nfatc1* expression and NFATc1 nuclear translocation.

**Figure 4.**
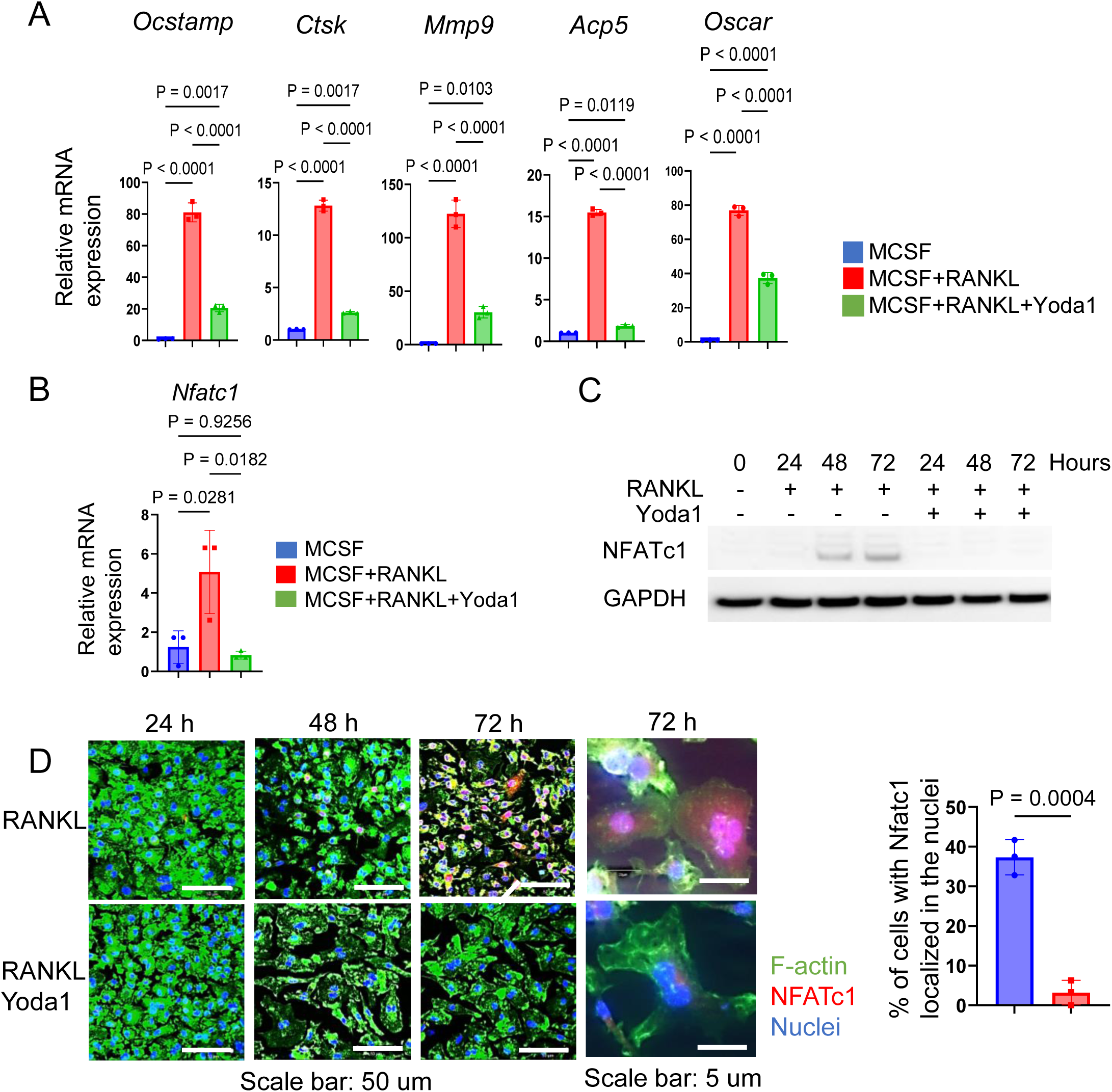
Pharmacological Piezo1 activation suppressed OC-related gene expression and NFATc1 expression in OCs. (A) Pre-OCs were treated (5 µM) or vehicle control. After 48 hours, *Ocstamp*, *Ctsk*, *Mmp9, Acp5* and *Oscar* were confirmed by qPCR. (B and C) NFATc1 mRNA and protein expression was determined from pre-OCs stimulated with or without Yoda1 at day 1, day 2 and day 3 by qPCR or Western blot analysis, respectively. GAPDH was loading control. (D) Immunofluorescence was employed to image the localization of NFATc1 in pre-OCs at day 1, day 2 and day 3. Cells with NFATc1 present in the nucleus were counted. Data represent the mean ± SD of three independent experiments.

### 5. Sensing of Shear stress by Piezo1 expressed on pre-OCs suppressed OC-genesis

RANK-positive mononuclear pre-OCs, which are derived from monocyte lineage cells, circulate through the vasculature and migrate to bone (54). Fluid shear stress influences the activities of immune cells, such as CD4 T cells, neutrophils and monocytes (55–57). As demonstrated above (Fig. 2D) and a previous report (58), Piezo1, but little, or no, Piezo2, is expressed by pre-OCs. Although one prior study regarding deletion of Piezo1 in osteoclast lineage cells did not identify a bone phenotype (59), the study did not evaluate mechanosignaling. To confirm the mechanosensitivity of Piezo1 expressed on pre-OCs, a loss-of-function assay employing RNA interference was performed. Reduced expression of Piezo1 mRNA and protein by Piezo1-specific siRNA (siPiezo1) relative to Control siRNA (siControl) was confirmed by qPCR and immunofluorescence (Fig. 5A). Yoda1-mediated induction of Ca^2+^ influx was diminished in siPiezo1-treated pre-OCs, but not in the siControl-treated group (Fig. 5B). The induction of Ca^2+^ influx in a 20 dyn/cm^2^ of shear flow-dependent manner was abrogated by treatment of pre-OCs with siPiezo1 (Fig. 5C). Also, mechanical stress generated by the shaker suppressed OC-genesis-related genes, including OCSTAMP, MMP9 and ACP5, expression, mature TRAP+ OC formation and pit formation, which was downregulated by the treatment of pre-OCs with siPiezo1 (Fig. 5D and E). These data indicated that Piezo1 expressed in pre-OCs is responsible for mechanical stress sensing by pre-OCs which, in turn, downregulates RANKL-induced OC-genesis.

**Figure 5.**
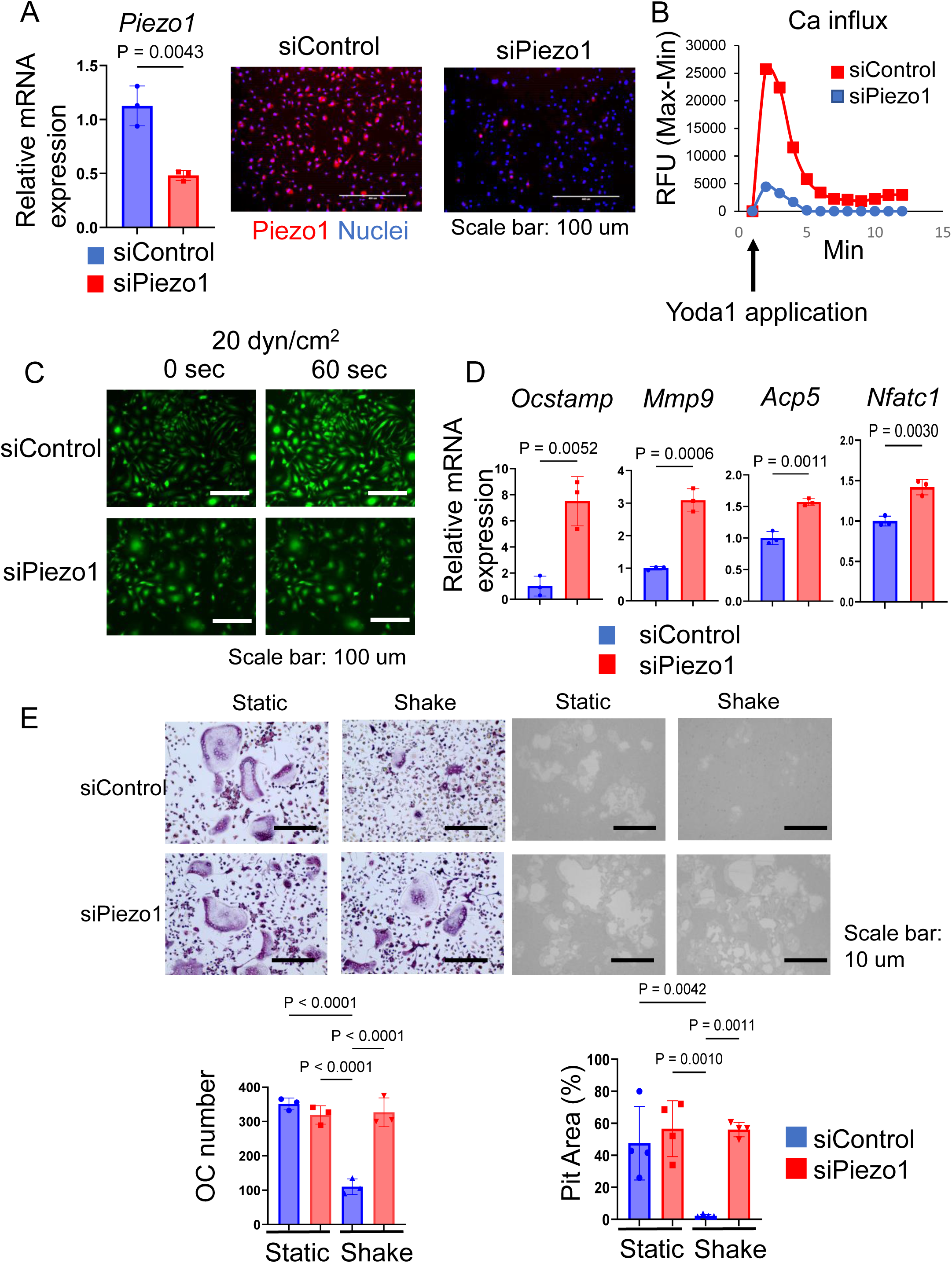
Piezo1 activated by shear stress downregulated OC-genesis. (A) To silence *Piezo1* expression, Pre-OCs were transfected with siRNA specific to Piezo1 (siPiezo1). siPiezo1-mediated silencing efficacy was evaluated by qPCR and immunofluorescence staining, in comparison to control siRNA treatment (siControl). (B) siPiezo1-mediated Piezo1 loss-of-function was determined by measuring Yoda1-enhanced Ca^2+^ influx. Fluo-8-treated pre-OCs were stimulated Yoda1, subsequently kinetic fluorescence intensity was measured every 1 min by plate reader. (C) Fluo-8-treated pre-OCs in µ-Slide I 0.4 Luer were stimulated 20 dyn/cm^2^ of shear stress generated by ibidi pump system, then time-lapse images were taken every 1 min. Representative image after 1 min shear flow exposure was displayed. (D) The siRNA-treated pre-OCs were stimulated with shear flow by shaking for 2 days. *Ocstamp, Mmp9*, *Acp5* and *Nfatc1* expression was analyzed by qPCR. (E) The number of TRAP-positive multinucleated OCs was measured, and measurement of pit area was performed. Data represent the mean ± SD of three independent experiments.

### 6. Piezo 1 activation by Yoda1 strongly suppressed Akt phosphorylation

Piezo1 is reported to provoke intracellular signaling activation in numerous cells (39, 60, 61). However, Piezo1-related signaling in OCs is still unknown. To address this question, phospho antibody array was performed to discover the specific signaling of Piezo1 in pre-OCs in *vitro*. A search of the Kyoto Encyclopedia of Genes and Genomes (KEGG) and Ingenuity® Pathway Analysis (IPA®) found PI3K/Akt signaling to be the most likely Piezo1 signaling pathway (Fig. 6A). Akt is a serine/threonine kinase that plays a critical role in cell survival, growth, and metabolism. It is well known that Akt is closely involved in OC-genesis (62, 63). Akt phosphorylation activates GSK3β that then evokes NFATc1 nuclear translocation during OC formation (64–66). As further confirmation, inhibition of PI3K/Akt signaling by LY294002 resulted in suppressed OC formation and Nfatc1 expression, suggesting that PI3K/Akt signaling is, indeed, involved in OC-genesis (Fig.S1B). In addition, Yoda1 application significantly reduced Akt phosphorylation and moderately inhibited ERK phosphorylation (Fig. 6B). Moreover, shear flow-mediated Akt dephosphorylation was regulated via Piezo1 (Fig. 6C and Fig. S6B). Therefore, Akt dephosphorylation induced by Piezo1 activation is associated with OC formation.

**Figure 6.**
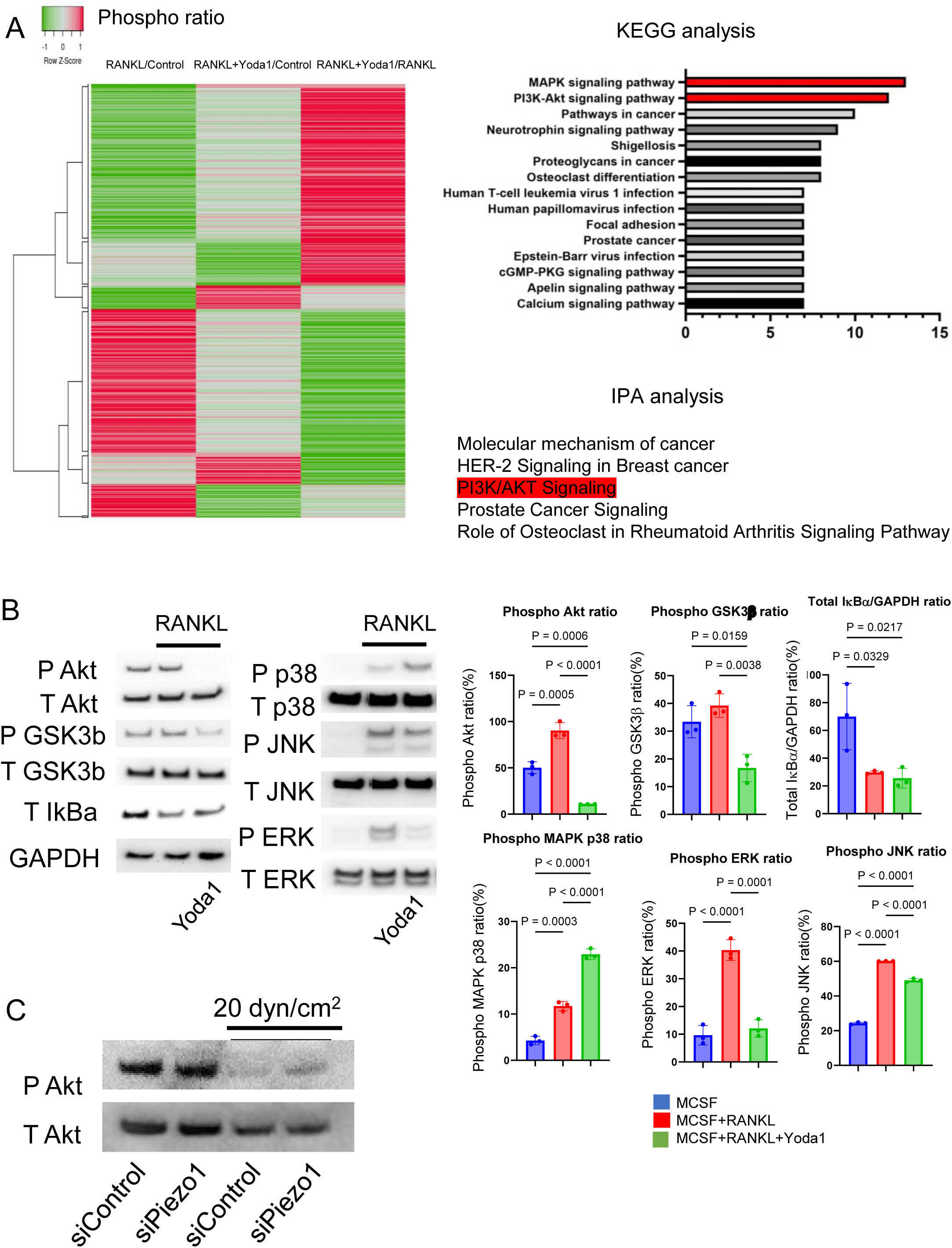
Piezo 1 activation by Yoda1 strongly suppressed Akt phosphorylation (A) Phospho Explorer Antibody Array was performed to determine Yoda1-mediated signaling pathway in pre-OCs. KEGG and IPA were used for bioinformatics analysis. ( (B) Pre-OCs were stimulated with RANKL (10 ng/ml) in the presence or absence of Yoda1 (5 uM) for 30 min to monitor protein phosphorylation, including Akt, GSK-3β, p38^MAPK^, ERK, JNK and NF-kB. siRNA-transfected OCs were cultured with shear flow, and samples were collected to monitor Akt phosphorylation by Western blot. Densitometric analysis was conducted using ImageJ software (Version 1.50). (C) Pre-OCs treated with siRNA for Piezo1 or negative control were stimulated with shear stress at 20 dyn/cm^2^, then western blotting was preformed to determine Akt phosphorylation. Representative band images are shown from three independent experiments. Data represent the mean ± SD of three independent experiments.

### 7. PP2A, not Calcineurin, is involved in Piezo1-induced Akt dephosphorylation in OCs

Akt is also regulated by protein phosphatase family, such as protein phosphatase 2A (PP2A) or protein phosphatase 2B (PP2B), known as calcineurin (67). We demonstrated that okadaic acid, a PP2A inhibitor, but not FK506, a calcineurin inhibitor, could counteract Yoda1-induced dephosphorylation of Akt (Fig. 7A and Fig. S6C). Subsequently, as a gain-of-function approach, DT-061, a PP2A activator (68), was employed to elucidate its functional role in OC-genesis. We demonstrated that DT-061, through its activation of PP2A, significantly suppressed TRAP+ OC formation, resorption pit formation, and Akt phosphorylation in RANKL-stimulated pre-OCs (Fig. S4A). RANKL-induced NFATc1 expression was also downregulated by treatment with the PP2A activator DT-061 (Fig. S4C). We found that Yoda 1 downregulated the induction of PP2A phosphorylation at *Tyr**^307^***, which elicits the catalytic activity of PP2A phosphatase (71,72) by RANKL-stimulated pre-OCs (Fig. 7B). It is noteworthy that the PP2A catalytic subunit (PP2Ac) is inactivated by single phosphorylation at *Tyr**^307^*** residue (69), whereas phosphorylation of *Tyr^127^* and *Tyr^284^* can activate PP2Ac (70). These findings indicate that activation of Piezo1 can suppress phosphorylation of PP2A at Tyr30 to increase phosphatase activity by PP2A which, in turn, suppresses the phosphorylation of Akt, i.e., Akt dephosphorylation, as well as NFATc1. RNAi-based silencing of PP2A mRNA expression (siPP2A), but not calcineurin (siCalcineurin), resulted in increasing Akt phosphorylation in Yoda1-treated OCs (Fig. 7C). Furthermore, treatment with siPP2A, but not siCalcineurin, prevented shear-stress-dependent suppression of RANKl-stimulated OC-genesis, otherwise activated by PP2A, including TRAP-positive OC formation, pit formation, and OC-genesis-related gene expression (Fig. 7D and E). The protein phosphatase 2A (PP2A) is well known to negatively regulate Akt activity. Collectively, these results suggested that Piezo1-mediated mechanosensing by pre-OCs suppresses RANKL-induced OC-genesis through cell signaling that involves the PP2A/Akt-dephosphorylation pathway toward the suppression of NFATc1, the master TF controlling OC-genesis.

**Figure 7.**
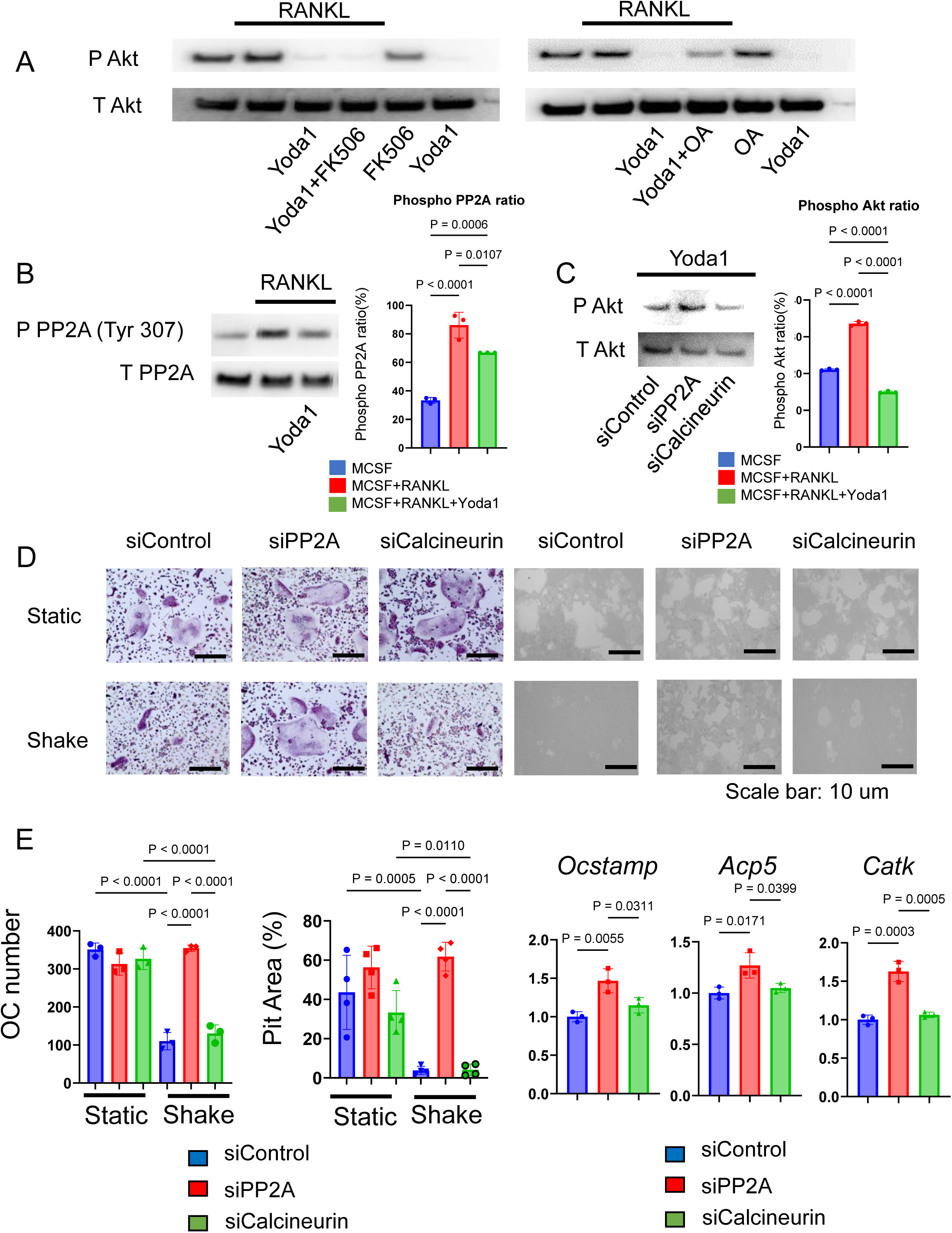
PP2A, not Calcineurin, is involved in Piezo1-induced Akt dephosphorylation in OCs. (A) Following preincubation with okadaic acid (protein phosphatase inhibitor) (250 nM) or FK506 (calcineurin inhibitor) (1 uM) for 1 hour, pre-OCs were incubated with Yoda1 (5 uM) for 30 min. Western blotting was performed to detect Akt phosphorylation. (B) Phosphorylation of the PP2A catalytic subunit at Tyr307 was visualized by Western blotting. Densitometric analysis was performed and data are shown. (C) Either siPP2A or siCalcineurin was employed to evaluate Yoda1-mediated Akt dephosphorylation by Western blotting. Densitometric analysis was performed and data are shown. (D and E) To determine the effect of PP2A or calcineurin on mechanical force downregulation of OC-genesis, pre-OCs were transfected with siPP2A, siCalcineurin or siControl, followed by TRAP staining, pit formation assay and qPCR for *Ocstamp, Acp5* and *Catk* expression. Data represent the mean ± SD of three independent experiments.

### 8. Yoda1 administration prevents osteoclastic bone resorption in a mouse model of ligature-induced periodontitis

Given the decreased shear stress and increased osteoclastic bone resorption observed in periodontitis, we examined the effect of systemic (i.p.) injection of Yoda1 in the mouse ligature-induced periodontitis model. Murine periodontitis was induced by the attachment of a silk ligature at the upper second molar, following previous reports (71–73). Systemically administered Yoda1 significantly suppressed bone resorption and ligature-induced TRAP-positive OC formation in alveolar bone compared to vehicle control (Fig. 8A). It also inhibited the mRNA expression of OCSTAMP, ACP5 and MMP9, but not RANKL mRNA (*Tnfsf11*) or osteoprotegerin mRNA (Tnfsf11b) (Fig. 8B). Furthermore, the number of phosphorylated-Akt-positive OCs was increased in mouse alveolar bone (Fig. 8C).

**Figure 8.**
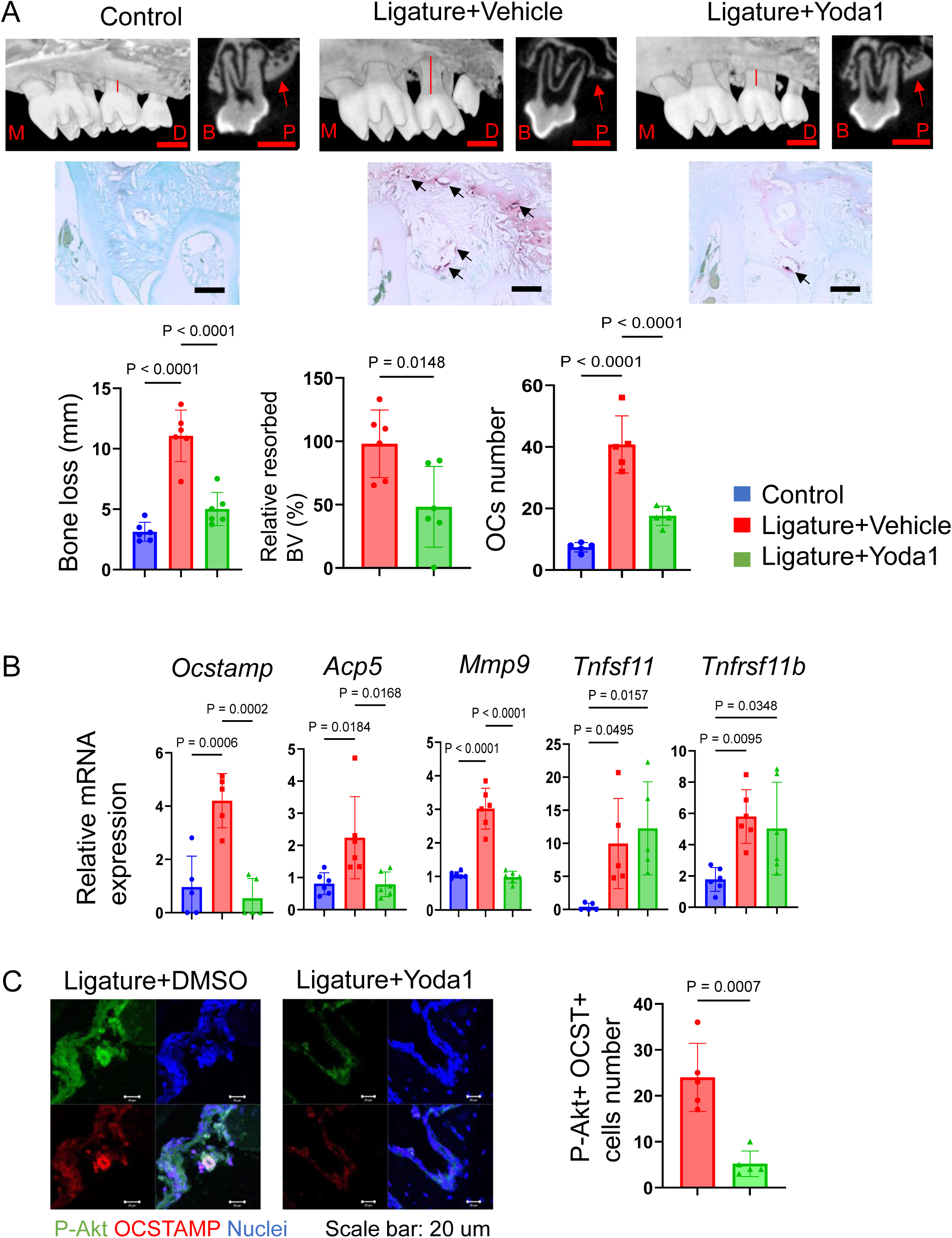
Yoda1 administration prevents osteoclastic bone resorption in murine ligature-induced periodontitis in mice. A 5-0 silk ligature was placed around murine maxillary second molar for 7 days to induce periodontitis. Yoda1 (0.4 mg/kg) or DMSO (as a vehicle control, 0.86 %) was administered systemically at day 0, 2, 4 and 6, respectively. (A) Bone resorption and TRAP-positive multinucleated OCs were evaluated by micro-CT analysis and TRAP staining. (B) Gingival tissue samples were harvested for qPCR to measure Ocstamp, Acp5, Mmp9, Tnfsf11 and Tnfsf11b. (C) Frozen tissue sections of mouse model of periodontitis were employed to image phospho-Akt and OC-STAMP double positive cells in mouse model of periodontitis. Number of double positive cells was counted. Results were presented as the means ± SD.

These results revealed that Piezo1 activation by Yoda1 directly reversed bone resorption in periodontitis by inhibiting mature OC formation.

## Discussion

Our findings suggested that Piezo1 negatively regulates OC-genesis in conjunction with the diminished expression of OC marker genes, including *Ocstamp*, *Acp5*, *Mmp9*, *Ctsk*, *Oscar* and *Nfatc1*, thereby suppressing bone resorption through its activation of negative-regulator PP2A that dephosphorylates Akt in the RANKL-induced TRAF6/PI3K/Akt/NFATc1 cell signaling axis. A mouse model of ligature-induced periodontitis demonstrated that Piezo1 activation by Yoda1 could downregulate pathogenically elevated OC-genesis in the alveolar bone of periodontally diseased tissue.

It was reported that murine arthritis-associated osteoclastogenic macrophages (AtoMs) comprise the CX_3_CR1^hi^ FoxM1^+^ pre-OCs-containing population in inflamed synovium and that they originate from circulating bone marrow cells (74).

Our group also demonstrated that locally produced macrophage migration inhibitory factor (MIF) in the inflammatory bone lytic site is engaged in the chemoattraction of circulating CXCR4^+^ pre-OCs to the inflammatory bone resorption site (22). These lines of evidence support that pre-OCs migrate from circulation to the inflammatory bone resorption site via chemotaxis, suggesting that pre-OCs, both during and after transvasation, i.e., pouring out of one vessel into another, in the inflammatory bone resorption site, are affected by the shear stress in local vasculature. Accordingly, owing to lower blood flow velocity in periodontitis (27), it is plausible that the diminished shear stress may affect the fate of pre-OCs differentiating into mature OCs. In support of this hypothesis, we observed a significant alteration in blood flow during murine ligature-induced periodontitis, characterized by reduced blood flow in periodontitis-affected tissue compared to healthy control tissue (Fig. 1). We found that pre-OCs express a functional Piezo1 mechanosensory ion channel (Fig. 2) as a kind of negative rescue factor by the imposition of shear force that otherwise downregulates OC formation via Piezo1 Ca^2+^ ion channel (Fig. 5). Thus, it was further hypothesized that Piezo1 may act as a major mechanoreceptor in circulating pre-OCs and that once activated, Piezo1 channels could initiate the PP2A-Akt signaling pathway to downmodulate the expression of genes associated with OC differentiation.

Piezo1 is a key mediator of mechanotransduction in bone cells, including osteoblasts, osteocytes and mesenchymal stem cells (75–77). It is involved in the differentiation of mesenchymal stem cells into osteoblasts or odontoblasts (77, 78), and is responsible for creating mechanical force and converting it into biochemical signals that regulate cellular responses. In response to mechanical stimuli, Piezo1 channels open, allowing the influx of Ca^2+^ into OBs; this Ca^2+^ influx then triggers a cascade of intracellular signaling events that ultimately lead to bone formation, including, as noted above, activation of ERK or, in our case, Akt cascade (79). Wang et al. reported that the Piezo1/YAP1/collagen pathway is associated with OB maturation *in vivo* and *in vitro* (59). Osteocytes also sense mechanical force through Piezo1, and intracellular signaling occurs in osteocytes through the Piezo1/Akt axis (76). Our data suggested that Piezo1 activation in OCs downregulated Akt signaling (Fig. 6 and 7), indicating that Piezo1 expressed in OC acts in a unique role compared to other bone cells. More specifically, based on our study and those of others, mechanosensing via Piezo1 not only promotes osteoblastic bone formation but also inhibits osteoclastic bone resorption through distinctly facilitated Piezo1-mediated cellular signaling pathways.

As previously noted, NFATc1 is a master TF controlling OC-genesis. Ligation of RANKL to RANK expressed on pre-OCs elicits cell signals involving the TRAF6/PI3K/Akt axis for induction of NFATc1 nuclear-translocation which, in turn, activates OC-genesis, (10, 80). However, Yoda1, the Piezo1 agonist, inhibited NFATc1 expression in pre-OCs stimulated with RANKL (Fig. 4B, C and D). Phospho Antibody Array (Fig. 6A) indicated that Akt plays a key regulatory function in Piezo1-elicited cell signals for OC-genesis. Indeed, the PI3K/Akt axis plays a crucial role in OC formation (65), whereas interaction of Akt-mediated activation of GSK-3β down-modulates OC formation via inhibition of nuclei translocation of NFATc1 (65, 81, 82). Therefore, we can conclude from the above findings that Piezo1-mediated cell signaling suppresses the Akt/NFATc1 axis.

PP2A and PP2B, also known as calcineurin, are protein phosphatases that dephosphorylate specific substrates and play important roles in cell signaling and regulation (83, 84). We demonstrated that Piezo1 activation in OCs promoted PP2A-mediated Akt dephosphorylation (Fig. 7A, B and C). Furthermore, we also determined that DT-061, a PP2A activator, exhibits a crucial preventive effect on OC-mediated bone resorption in murine periodontitis (Fig. S4B). Myung et al. reported that PP2A inactivation promotes OC-genesis (85). Hyun-Jung et al. also indicated that Dauricine, an isoquinoline alkaloid, decreases OC formation via activation of PP2A (86). These reports and our results strongly suggest that activation of PP2A in OCs can negatively control their differentiation. Furthermore, we discovered, for the first time, that activation of Piezo1 negatively regulates OC-genesis via the PP2A/Akt axis. On the other hand, calcineurin (PP2B) is a Ca^2+^- and calmodulin-dependent serine/threonine protein phosphatase (87). Calcineurin inhibition by FK506 or siRNA was ineffective in dephosphorylation of Akt and failed to abrogate shear stress-mediated suppression of OC formation (Fig. 7 A and D). Indeed, chemical-based inhibition of calcineurin results in the induction of osteoblastic bone formation (91, 92), but suppression of OC formation (93, 94). Therefore, we concluded that Piezo1-mediated dephosphorylation of Akt depends on PP2A, not calcineurin, in OCs. Moreover, since Piezo1 activation causes Ca^2+^ influx and Ca^2+^ -dependent cellular activity (95, 96), we concluded that Piezo1 activation in OCs may be involved in Ca^2+^-dependent PP2A activation. On the other hand, Ca^2+^ binding is reported to enhance the interaction between the B″/PR72 family, which belongs to the PP2A subunit, and the PP2A core enzyme, in turn promoting phosphatase activity (97, 98). Further studies are needed to adequately address the mechanisms influencing Piezo1-related OC formation associated with the Ca^2+^-dependent B"/PR72 family.

Ca^2+^ influx is strongly associated with OC-genesis. RANKL/RANK binding allows Ca^2+^ influx, following NFATc1 activation (10). RANKL-mediated OC genesis requires a costimulatory signal characterized by Ca^2+^ influx from ITAM receptors, such as OSCAR and TREM2, triggered by type 3 collagen (11, 12). However, both RANKL and type 3 collagen did not induce Ca^2+^ influx in pre-OCs (Fig. S3). Ionomycin, a Ca^2+^ ionophore, is reported to induce Ca^2+^ influx and OC-genesis (99). Instead, however, ionomycin administration at previously reported concentration (500 nM) increased Ca^2+^ influx, but suppressed OC-genesis (Fig. S5 A, B and C). Thus, it was clear that intracellular calcium influx is not necessarily a positive regulator of OC-genesis.

Ligature-induced periodontitis in mice is a well-established model of periodontitis as published in our previous reports (73, 100, 101). Here, we demonstrated that systemic Yoda1 application in mice significantly prevented murine periodontal bone loss induced by placement of ligature (Fig. 8A). Yoda1 is widely used as a specific pharmacological activator of Piezo1 and has applications in the analysis of the bioactivity of Piezo1 in various cells (46, 102, 103). For example, Yoda1 administration in mice enhanced microglial phagocytosis resulting in Aβ clearance in Alzheimer’s disease (104). Yoda1 administration did not alter the body weight of mice; instead, it increased cortical thickness and cancellous bone mass in the distal femur of mice (105). Our results indicated that Yoda1 alone did not affect bone resorption in control without ligature group (data not shown). In addition to significantly improving bone loss in the mouse model of ligature-induced periodontitis, Yoda1 suppressed gene markers of OC-genesis, including *Ocstamp*, *Acp5* and *Mmp9*, but not *Tnfsf11* and *Tnfsf11b* (Fig. 8 B), and phosphorylated Akt-positive OCs at the bone surface of murine periodontitis (Fig. 8 C). These results suggest that Yoda1 directly suppressed ligature-induced OC formation *in vivo*.

In summary, we have identified that pre-OCs express functional Piezo1 and that mechanical and chemical activation of Piezo1 expressed on pre-OCs downregulates OC-genesis through dephosphorylation of Akt by PP2A, which, in turn, suppresses the expression of NFATc1, a master TF for RANKL-induced OC-genesis (Fig. 9). Furthermore, stimulating Piezo1 via systemic administration with Yoda1 appeared to abrogate the diminished mechanostress in the inflamed periodontium in the mouse model of periodontitis, resulting in the inhibition of local bone loss mediated by osteoclasts. Therapeutically breaking through this feedforward mechanism provides a reasonable target in the search for novel treatments for periodontitis.

**Figure 9.**
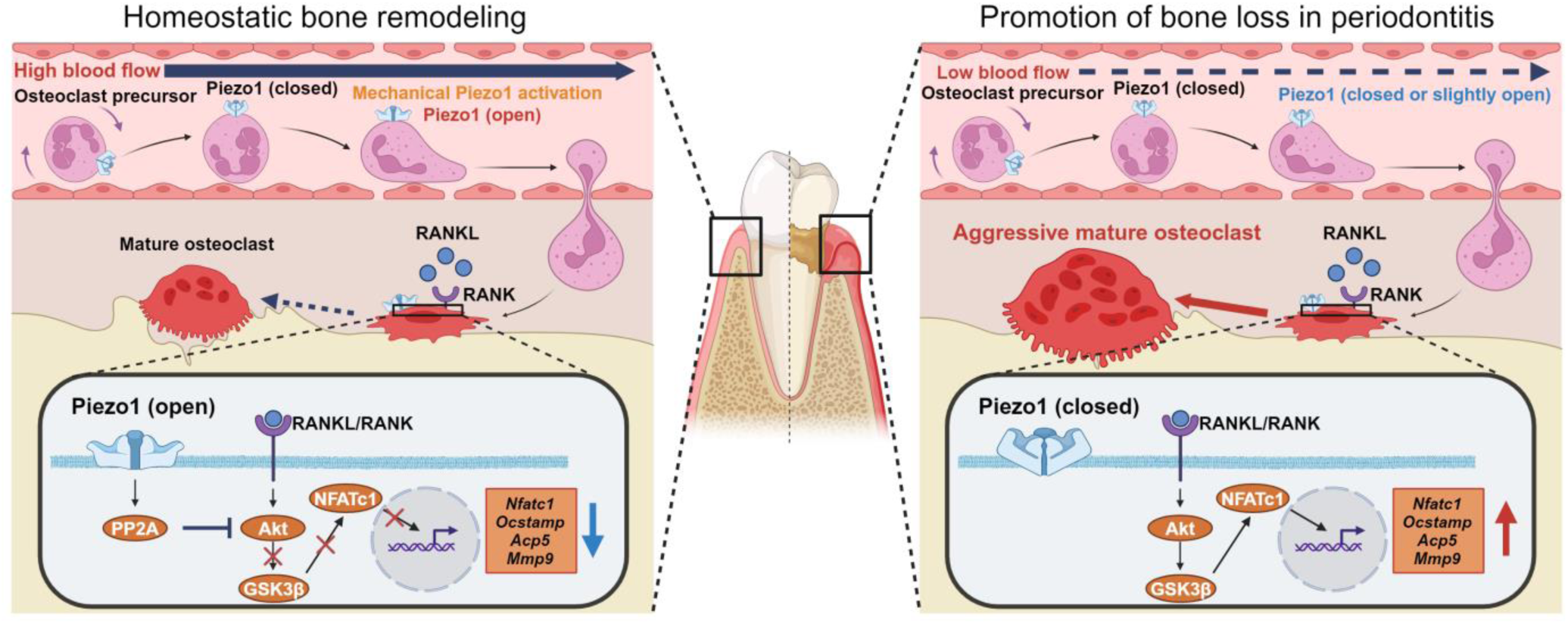
The role of Piezo1 expressed on OCprecs in periodontitis. In healthy periodontal tissue, OCprecs receive high blood flow in the vasculature, which enhances Piezo1 activation and continues PP2A/Akt signaling, resulting in regulated OC differentiation. However, low blood flow in diseased vasculature results in dysregulation of Piezo1 activation, causing excess OC differentiation to induce bone loss in alveolar bone.

## Materials and Methods

Materials and Methods are described in Supporting information.

## Acknowledgments

The schematic figures were created with BioRender.com. This work was supported by NSU PFRDG grant 334930, NIH NIDCR grants DE025255, DE027851, DE028715, DE029709, DE032907, as well as NIGMS grant GM150469. S.S. was supported by JSPS Overseas Research Fellowships.

## Author Contributions

S.S. and T.K. designed research; S.S., X.H., S.K. and T.K., secured research operation funds; S.S., S.N., M.R.Q., A.H., M.R.P., M.O., M.S., M.S., and A.B. performed research; S.S., S.N., M.R.Q., M.S., D.M., M.C., Y.Y., J.L.C., M.H., P.H., and X.H., analyzed data; S.S., S.V. and T.K. wrote the paper.

## Classification

Major; Biological Sciences, and Minor; Cell Biology

