## Supplemental file for "Mechanosensing by Piezo1 regulates osteoclast differentiation via PP2A-Akt axis in periodontitis"

Supporting information for **Mechanosensing by Piezo1 regulates osteoclast differentiation via PP2A-Akt axis in periodontitis**

Satoru Shindo^1^, Shin Nakamura^1^, Mohamad Rawas-Qalaji^1^, Alireza Heidari^1^, Maria Rita Pastore^1^, Motoki Okamoto^1^, Maiko Suzuki^1^, Manuel Salinas^2^, Dmitriy Minond^3^, Alexander Bontempo^1^, Mark Cayabyab^1,8^, Yingzi Yang^4^, Janet L Crane^5^, Maria Hernandez^6^, Saynur Vardar^6^, Patrick Hardigan^8^, Xiaozhe Han^1, 8^, Steven Kaltman^7^, Toshihisa Kawai^1, 8^ *

* Correspondence; Toshihisa Kawai

Department of Oral Science and Translational Research, College of Dental Medicine, Nova Southeastern University, Fort Lauderdale, FL33328, USA Tel: 954-262-1282

This PDF file includes:

Materials and Methods

Figure S1 to S6

**Materials and Methods**

1. Animals

C57BL/6 mice (8 to 10 weeks old female mice) were kept in a conventional room with a 12-h light-dark cycle at constant temperature. The experimental procedures employed in this study were approved by the NSU IACUC (Protocol #TK7).

2. Cell Culture

Bone marrow-derived mononuclear cells (BMMCs) were collected from tibias and femurs of C57BL/6 mice and were cultured with minimum essential medium-α (MEM-α) supplemented with 10% fetal bovine serum (FBS), streptomycin (100 µg/mL), penicillin (100 U/mL), L-glutamine (292 μg/mL) and M-CSF (25 ng/mL; BioLegend, San Diego, CA, USA) at 37 °C in humidified air with 5% CO_2_ for 3 days to obtain pre-OCs. Subsequently, pre-OCs were differentiated to OCs using RANKL (10 ng/mL; BioLegend). Human Peripheral Blood Mononuclear Cells (PBMCs) were purchased from STEMCELL Technologies (Vancouver, Canada). PBMCs were stimulated with M-CSF (30 ng/mL; BioLegend) at 37 °C in humidified air with 5% CO_2_ for 3 days. Human pre-OCs were differentiated to OCs using RANKL (100 ng/mL; BioLegend). Yoda1 (Tocris), GsMTx4 (Tocris), LY294002 (Cell Signaling Technology), FK506 (Tocris), Ocadaic acid (R and D system) and DT-061 (Tocris) were administered with validated concentration. Aa a vehicle control, same concentration of Dimethylsulfoxide (DMSO, Sigma) was used.

3. Cell culture under flow

Shear stress was generated by shaker (Thermo Fisher Scientific) or ibidi pump system (ibidi, Fitchburg, WI). Pre-OCs were seeded in wells of a 24-well plate (1x10^6^ cells/well) and then stimulated with shear stress by shaking (15°, 30 rpm). On the other hand, for the ibidi pump system, pre-OCs were cultured in µ-Slide I 0.4 Luer (3x10^5^ cells/well). Cells were cultured with shear stress at 5 dyn/cm^2^ or 20 dyn/cm^2^.

4. Tartrate-Resistant Acid Phosphatase (TRAP) Staining

TRAP staining was performed with a TRAP staining kit (Sigma-Aldrich) according to the manufacturer’s protocol. Briefly, OCs were fixed by a citrate (0.38 mol/L)/acetone solution for 30 seconds at room temperature. Cells were stained with a TRAP staining solution (L(+)-tartrate buffer, 0.67 mol/L; acetate buffer, 2.5 mol/L; Naphthol AS-BI phosphoric acid, 12.5 mg/mL; and Fast Garnet GBC base, 7.0 mg/mL) for 10 min at 37 °C in the dark. OCs with ≥ 3 nuclei were determined as multinucleated OCs.

5. Pit Formation Assay

A plate coated with calcium phosphate was prepared according to previous reports (106, 107). Briefly, 0.12 M Na_2_HPO_4_ and 0.2 M CaCl_2_ (50 mM Tris-HCl, pH 7.4) were mixed at 37 °C. The calcium phosphate slurry was washed with sterile water and then applied into wells of a culture plate and dried at 37 °C overnight. BMMCs or RAW264.7 cells (3.0 × 10^5^ cells/well for 96-well plate or 1× 10^6^ cells/well for 24- well plate) were then seeded in wells of calcium phosphate-coated plates, respectively. After treating cells for 6 days, the plates were washed with 10% sodium hypochlorite for 10 min and then dried overnight. The pit areas were microscopically imaged (Evos Cell Imaging System, Thermo Fisher Scientific). Images were analyzed using ImageJ software (version 1.50).

6. Measurement of intracellular Ca^2+^ concentration

Cells (4×10^4^ cells/well) were seeded in wells of a black wall/clear bottom plate. Fluo-8 No Wash Calcium Assay Kit (AAT Bioquest, Pleasanton, CA) was employed to measure Ca^2+^ influx in accordance with the manufacturer’s protocol. Briefly, Fluo-8 NW and 0.04% Pluronic™ F-127 in HHBS buffer were added for 30 min at 37 °C, followed by 30 min at room temperature. After Yoda1 treatment, fluorescence was alternately excited at 490 nm and emission at 525 nm measured every 10 sec using a FilterMax F5 Microplate Reader (Molecular Devices, San Jose, CA).

Ca^2+^ influx was also analyzed under flow conditions with the BioFlux One system (Fluxion Biosciences, Oakland, Ca) or ibidi pump system (ibidi). For the BioFlux One system, a 48-well microfluidic plate (Fluxion Biosciences) was first coated for 1 h at room temperature with rat tail type 1 collagen (50 μg/ml; Thermo Fisher Scientific) in 0.2% acetic acid. Before using the plate, microfluidic channels were washed with PBS, followed by the introduction of cells in the channels. After 24 h, the Fluo-8 No Wash Calcium Assay Kit was employed to image Ca^2+^ influx. The assay was performed with a wall shear stress at 20 dyn/cm^2^. Fluorescence intensity was measured and analyzed by the BioFlux One system (Fluxion Biosciences). For ibidi pump system, pre-OCs were seeded in µ-Slide I 0.4 Luer (ibidi) at 3x10^5^ cells/plate. The assay was performed with a wall shear stress at 20 dyn/cm^2^. Fluorescence intensity was measured and analyzed by the EVOS (Thermo Fisher Scientific).

7. siRNA Transfection for Knockdown of Piezo1.

BMMCs were seeded in wells of a 96-well plate (3×10^5^ cells/well) or 24-well plate (1×10^6^ cells/well). After 24 h, BMMCs were maintained in Opti-MEM™ Reduced Serum Medium (Thermo Fisher Scientific) and transfected with 50 nM Piezo1-specific siRNA (107969, Thermo Fisher Scientific; siPiezo1), PP2A-specific siRNA (152168, Thermo Fisher Scientific; siPP2A), PP2B-specific siRNA (162266, Thermo Fisher Scientific; siPP2B) or negative control siRNA (AM4611, Thermo Fisher Scientific; siCTL) using Lipofectamine™ RNAiMAX Transfection Reagent (Thermo Fisher Scientific) according to the manufacturer's instructions. 48 h post- transfection, BMMCs were used for indicated experiments.

8. Quantitative Polymerase Chain Reaction (qPCR)

Total RNA was extracted from pre-OCs using the PureLink RNA Mini Kit (Thermo Fisher Scientific), following the manufacturer’s protocol. The first strand cDNA was assembled from 100 ng of sample RNA using a Verso cDNA Synthesis Kit (Thermo Fisher Scientific). Amplification reactions were performed by Taqman Fast Advanced Master Mix (Thermo Fisher Scientific). The resultant cDNA was amplified by specific probes (Thermo Fisher Scientific) for *Gapdh* (Mm99999915_g1), *Piezo1* (Mm01241549_m1), *Piezo2* (Mm01265861_m1), *Trpa1* (Mm01227437_m1), *Trpv4* (Mm00499025_m1), *Stoml3* (Mm01289590_m1), *Kcnk10* (Hs01026663_m1), *Kcnk4* (Mm00434626_m1), *Kcnk1* (Mm00434624_m1), *Ocstamp* (Mm00512445_m1), *Mmp9* (Mm00442991_m1), *Ctsk* (Mm00484039_m1), *Acp5* (Mm00475698_m1), Oscar (Mm01338227_g1), *Nfatc1* (Mm00438670_m1), *Tnf* (Mm00443258_m1), *Il1b* (Mm00434228_m1), *Tnfsf11* (Mm00441906_m1) and *Tnfsf11b* (Mm00435454_m1) on a QuantStudio™ 3 (Thermo Fisher Scientific). The ratios of mRNA levels to those of the control gene were calculated using the ΔCt method (2^−ΔΔCt^).

9. PCR Array

After extraction of total RNA from cells and synthesis of cDNA described above, the resultant cDNA was tested using Taqman Fast Advanced Master Mix (Thermo Fisher Scientific) and TaqMan® Array Mouse Osteogenesis (Thermo Fisher Scientific) according to the manufacturer’s instruction. An integrated web-based software package was used for data analysis (https://www.thermofisher.com/account-center/simplified-username.html).

10. Phospho Antibody Array

The Phospho Explorer Antibody Array was used according to the manufacturer’s instruction (Full Moon BioSystems, Sunnyvale, CA) to profile the levels of phosphorylated proteins. Briefly, cell lysate from BMMCs was collected and quantified by BCA protein assay kit (Thermo Fisher Scientific). Microarray slides were blocked, and proteins were labeled using biotin and then coupled to slides. Slides were washed, and Cy3-streptavidin was added to bind biotin. Fluorescence intensity was measured and analyzed by the manufacturer (Full Moon BioSystems). The clustering of target proteins and signaling pathways was assessed using Kyoto Encyclopedia of Genes and Genomes (KEGG) and Ingenuity Pathway Analysis (IPA) (Qiagen, Germantown, MD).

11. Western blotting

After incubation for various times, pre-OCs were lysed by incubation on ice for 30 min with RIPA buffer (Thermo Fisher Scientific) supplemented with a protease inhibitor cocktail (Sigma-Aldrich). Protein concentration of the resultant lysates was measured with the BCA Protein Assay Kit (Thermo Fisher Scientific). Fifteen μg of sample per lane were loaded onto a 4–12% sodium dodecyl sulfate–polyacrylamide gel electrophoresis (SDS-PAGE) gel (Thermo Fisher Scientific). Proteins separated in the SDS-PAGE gel were electrotransferred to a polyvinylidene difluoride membrane. The detection of specific proteins in pre-OCs was assessed using anti-phospho-Akt rabbit mAb (1:1000; Cell Signaling Technology, Danvers, MA), anti-Akt rabbit mAb (1:1000; Cell Signaling Technology), anti-phospho-p38 MAPKs rabbit mAb (1:1000; Cell Signaling Technology), anti-p38 MAPKs rabbit mAb (1:1000; Cell Signaling Technology), anti-phospho-ERK rabbit mAb (1:2000; Cell Signaling Technology), anti-ERK rabbit mAb (1:2000; Cell Signaling Technology), anti-phospho-JNK rabbit mAb (1:1000; Cell Signaling Technology), anti-JNK rabbit mAb (1:1000; Cell Signaling Technology), anti-IkBa rabbit mAb (1:1000; Cell Signaling Technology), anti-phospho-PP2A mouse mAb (1:1000; Santa Cruz Biotechnology, Dallas, TX), anti-PP2A mouse mAb (1:1000; Santa Cruz Biotechnology) or an anti-GAPDH rabbit mAb (Cell Signaling Technology). Protein bands that reacted with the respective antibody were visualized by incubation with an HRP-conjugated rabbit or mouse secondary antibody (Cell Signaling Technology), followed by detection using ECL Western Blotting Substrate (Thermo Fisher Scientific). Densitometric analysis was performed using ImageJ software (Version 1.50).

12. Immunocytochemistry

Pre-OCs cultured on Millicell EZ SLIDE (Sigma) were fixed with 4% paraformaldehyde at room temperature for 10 min, permeabilized with 2% Triton X-100, and blocked with 5% BSA in PBS for 1 h. Cells were incubated with anti-Piezo1 antibody conjugated with Alexa Fluor® 594 (1 ug/ml, Novus, Centennial, CO), isotype control antibody conjugated with Alexa 594 (1 ug/ml, Novus), anti-NFATc1 antibody (1 ug/ml, Santa Cruz Biotechnology) or isotype control antibody (1 ug/ml, Bio X Cell, Lebanon, NH) at 4 °C overnight. Cy3-conjugated goat anti-mouse IgG secondary antibody (1:100, Jackson ImmunoResearch, West Grove, PA) was applied for 1 hour at room temperature, followed CellMask™ Green Actin Tracking Stain (Thermo Fisher Scientific) and Fluoromount-G containing DAPI (Thermo Fisher Scientific). Immunofluorescence was imaged with a Zeiss LSM880 confocal microscope (Carl Zeiss, Jena, Germany).

13. Flow Cytometry

Pre-OCs (1×10^6^ cells) were stained with anti-Piezo1 antibody conjugated with Alexa 594 (Novus) or isotype control antibody conjugated with Alexa 594 (Novus) for 30 min on ice and then analyzed by a BD LSRFortessa (BD Biosciences, Franklin Lakes, NJ, USA) along with FlowJo v10 software (BD Biosciences).

14. Mouse Model of Periodontal Disease

To induce periodontal disease in C57BL6/J mice (6–8 weeks old; male n = 5/group), the maxillary second molar was attached with a 5-0 silk ligature, following the previously published protocol (71, 73). Yoda1 (0.4 mg/kg), DT-061 (0.4 mg/kg) or DMSO (0.86 %) was diluted in PBS with 5% ethanol, followed by injection through the intraperitoneal route every 2 days (day 0, 2, 4 and 6). After 7 days, mice were euthanized for postmortem analyses.

15. Monitoring blood perfusion unit (BPU)

Real time BPU was monitored by Laser Doppler Flowmetry (OxyFlo pro, Oxford Optronix, UK). A fine needle-type blood flow probe (Diameter: 0.5 mm, Oxford Optronix) was attached to the mesial or distal surface of palatal gingiva, respectively (Fig. 1). Real time data were captured and analyzed by Labchart 8 software (ADInstruments, Colorado Springs, CO).

16. Measurement of Alveolar Bone Resorption

Resected maxillae were mechanically removed and exposed to 3% hydrogen peroxide overnight to remove all soft tissue. To evaluate periodontal bone resorption, distances from the cement–enamel junction to the alveolar bone crest on the buccal side of each root were measured for the maxillary second molar under the microscope (Zeiss).

17. Histological Analysis

Murine maxillary bones were fixed in 4% paraformaldehyde overnight at 4 °C before decalcification in 10% EDTA at 4 °C for 2 weeks. Tissues were embedded in an OCT compound (Sakura Finetek USA, Torrance, CA, USA) overnight at −20 °C and cut into 8 μm sections with a cryostat (Leica Biosystems, Deer Park, IL). TRAP staining of decalcified periodontal tissue was performed using an Acid Phosphatase Leukocyte (TRAP) Kit (Sigma-Aldrich), as described above, followed by nuclear counterstaining with methyl green. Sections were imaged with an EVOS XL Core microscope (Thermo Fisher Scientific). For immunofluorescence-based detection of OC-STAMP and phospho-Akt, the sections were reacted with anti-OC-STAMP rabbit pAb (1:200; Sigma-Aldrich) or anti-phospho-Akt rabbit mAb (1:200; Cell Signaling Technology) as the primary antibody at 4 °C overnight. Cy3-conjugated anti-rabbit IgG FC goat pAb (1:200; Jackson ImmunoResearch) was used as a secondary antibody. The stained sections were mounted with Fluoromount-G containing DAPI (Thermo Fisher Scientific). Immunofluorescence was observed with a Zeiss LSM880 confocal microscope (Carl Zeiss, Jena, Germany).

18. Micro-CT analysis

Mouse maxillary alveolar bone was fixed in 4% phosphate-buffered paraformaldehyde and stored at 4˚C for 16 hours. Micro-CT images were obtained with a microfocus x-ray CT imaging system (Skyscan 1176, Bruker, Billerica, MA), using the following settings: acceleration voltage, 50 kV; current, 500 µA; voxel size, 18 µm/pixel; matrix size, 2,000 × 1,336. Images were reconstructed with NRecon software, version 1.7.0.3 (Bruker), and images of both ligature side and control untreated side were acquired. As regions of interest, 50 sliced images coronally from the contact point between the maxillary first molar and maxillary second molar were evaluated. Bone volume (BV) of the whole palatal alveolar bone, including the ipsilateral hard palate, was measured using three-dimensional (3D) analysis CTAn software, version v.1.18 (Bruker). 3-dimensional images were obtained using CTVox software, version 3.2.0 (Bruker).

19. Statistical Analysis

Statistical analyses were performed by one-way ANOVA and Tukey HSD to compare differences among multiple groups and Student’s t-test for comparisons between two groups. All statistical analyses were performed using GraphPad Prism, version 10.0.1 (GraphPad Software, Inc., La Jolla, California, USA). Statistical significance was considered to be at p < 0.05. All data were expressed as the mean ± SD.

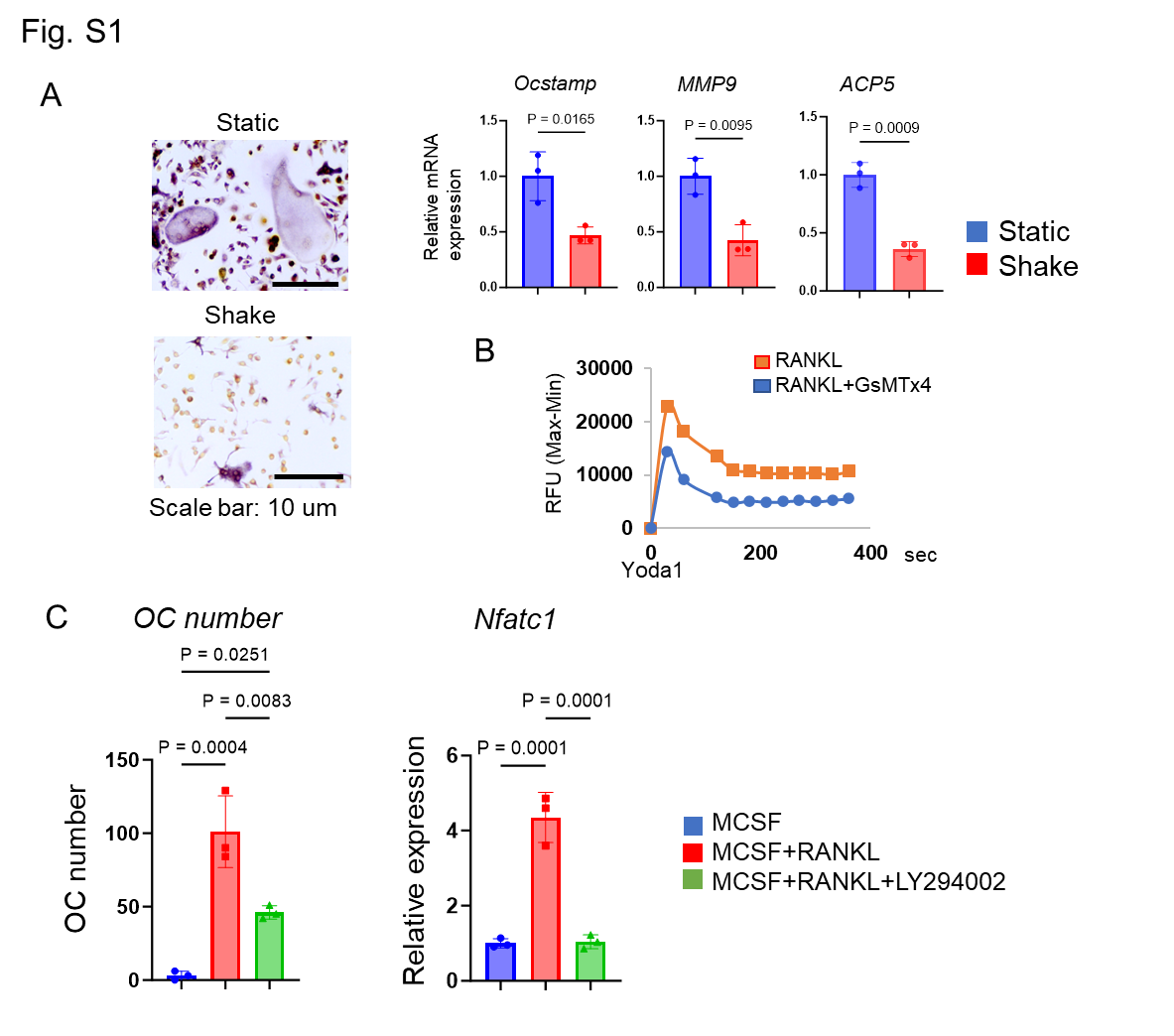

Figure S1. Shear stress generated by shaking inhibits OC-genesis, and GsMT-x4, Piezo1 inhibitor, suppresses Ca^2+^ influx mediated by Yoda1.

(A) Mechanical loading by rocker (15°, 30 rpm) was given pre-OCs to determine OC-genesis. (B) Yoda1-induced Ca^2+^ influx in pre-OCs with or without GsMT-x4 (1 uM) was measured. (C) B) Using LY294002 as an Akt inhibitor, TRAP staining and Nfatc1 expression were each analyzed to confirm the importance of Akt signaling in OC-genesis. Data represent the mean ± SD of three independent experiments.

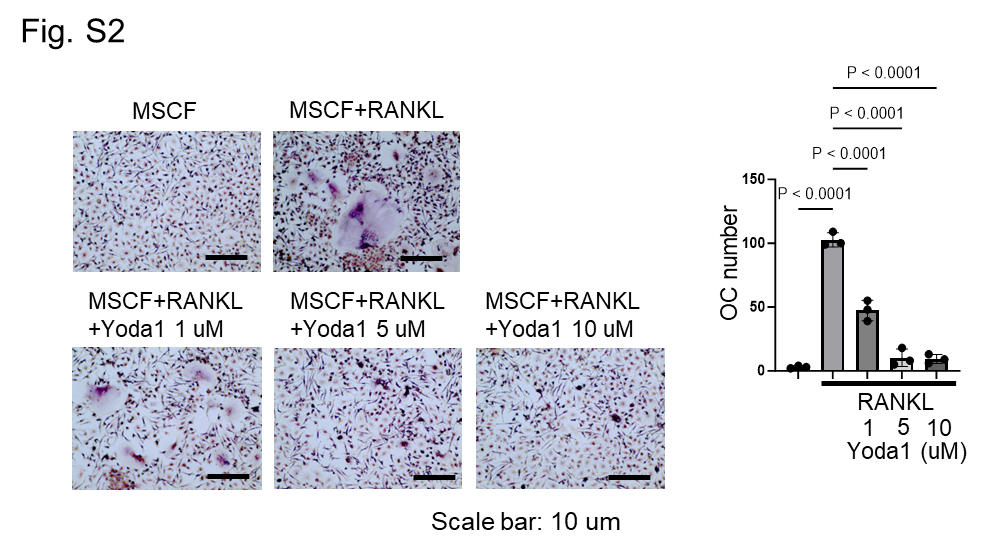

Figure S2. Piezo1 activation also inhibits human OC-genesis.

Yoda1 (1, 5, 10 uM) was applied RANKL-primed human pre-OC. TRAP positive multinucleated OCs were counted. Data represent the mean ± SD of three independent experiments. Results were presented as the means ± SD.

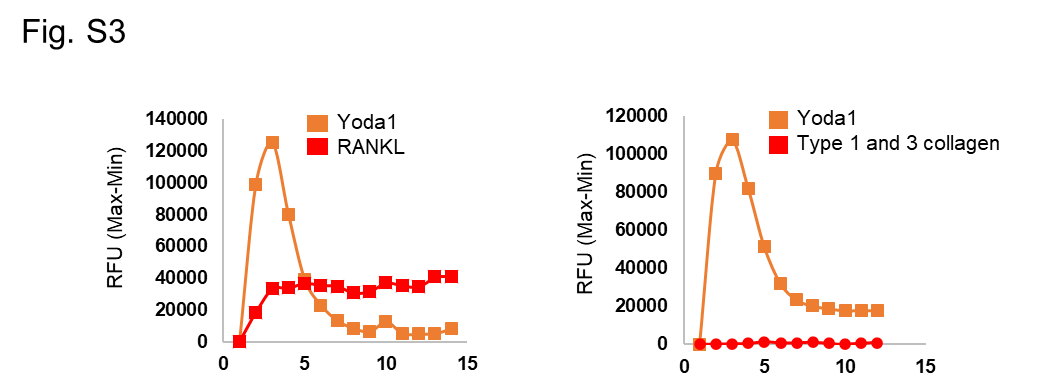

Figure S3. RANKL or type 1 and 3 collagen do not induce Ca^2+^ influx in OCs.

Pre-OCs were labeled with Fluo-8 NW, then Ca^2+^ influx was detected by a FilterMax F5 Microplate Reader.

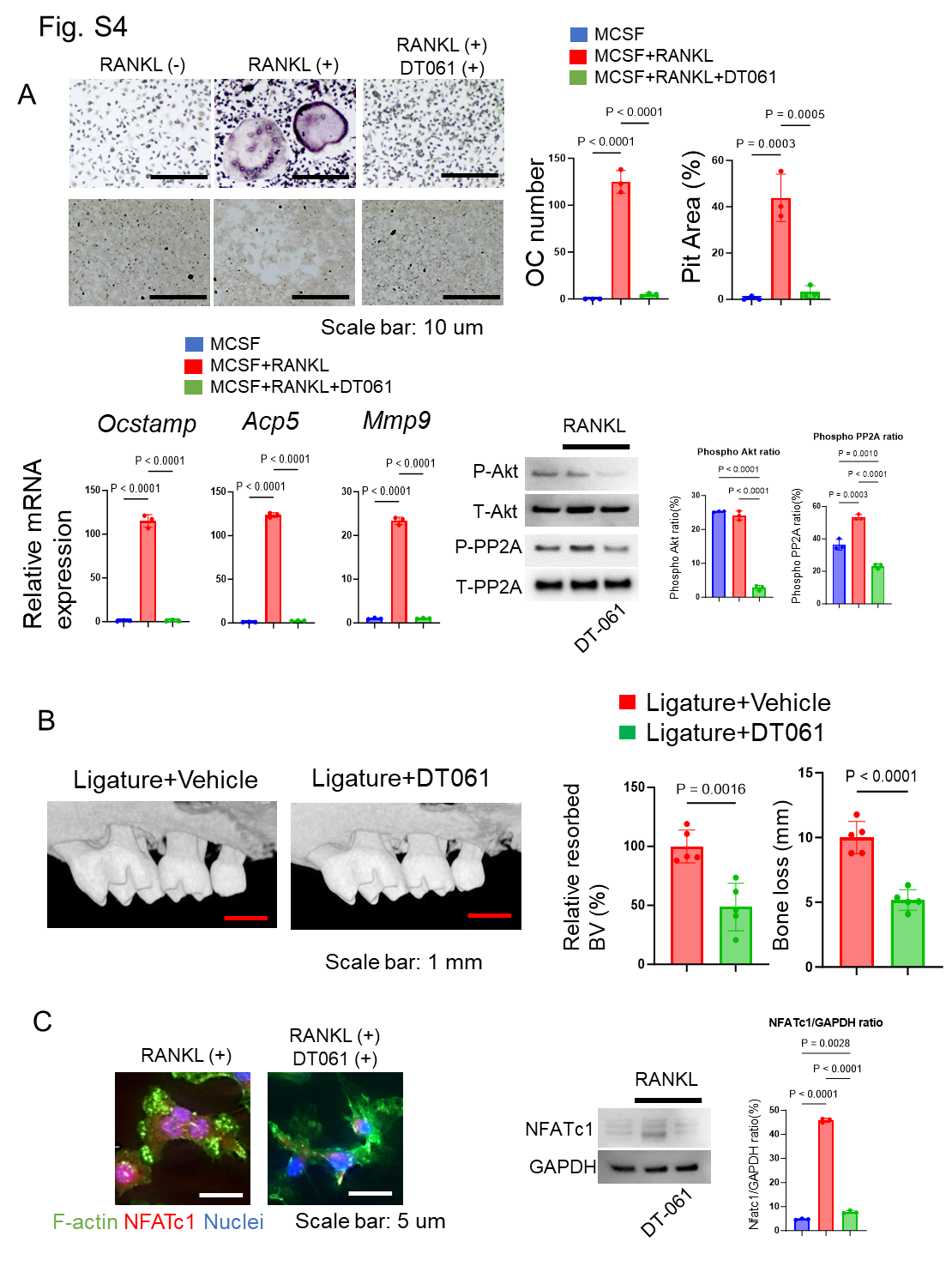

Figure S4. DT-061, PP2A activator, attenuates OC-genesis via Akt signaling in periodontitis.

(A) DT-061 (5 uM) or vehicle control were added pre-OCs to analyze OC-related gene expression including Ocstamp, Acp5 and Mmp9, along with RANKL-mediated Akt phosphorylation. (B) DT-061 (0.4 mg/kg) or vehicle control (DMSO. 0.86 %) was locally injected into the periodontitis area of mice induced by silk ligation. (C) DT-061 (5 uM) or vehicle control-mediated NFATc1 protein expression were imaged. Results were presented as the means ± SD.

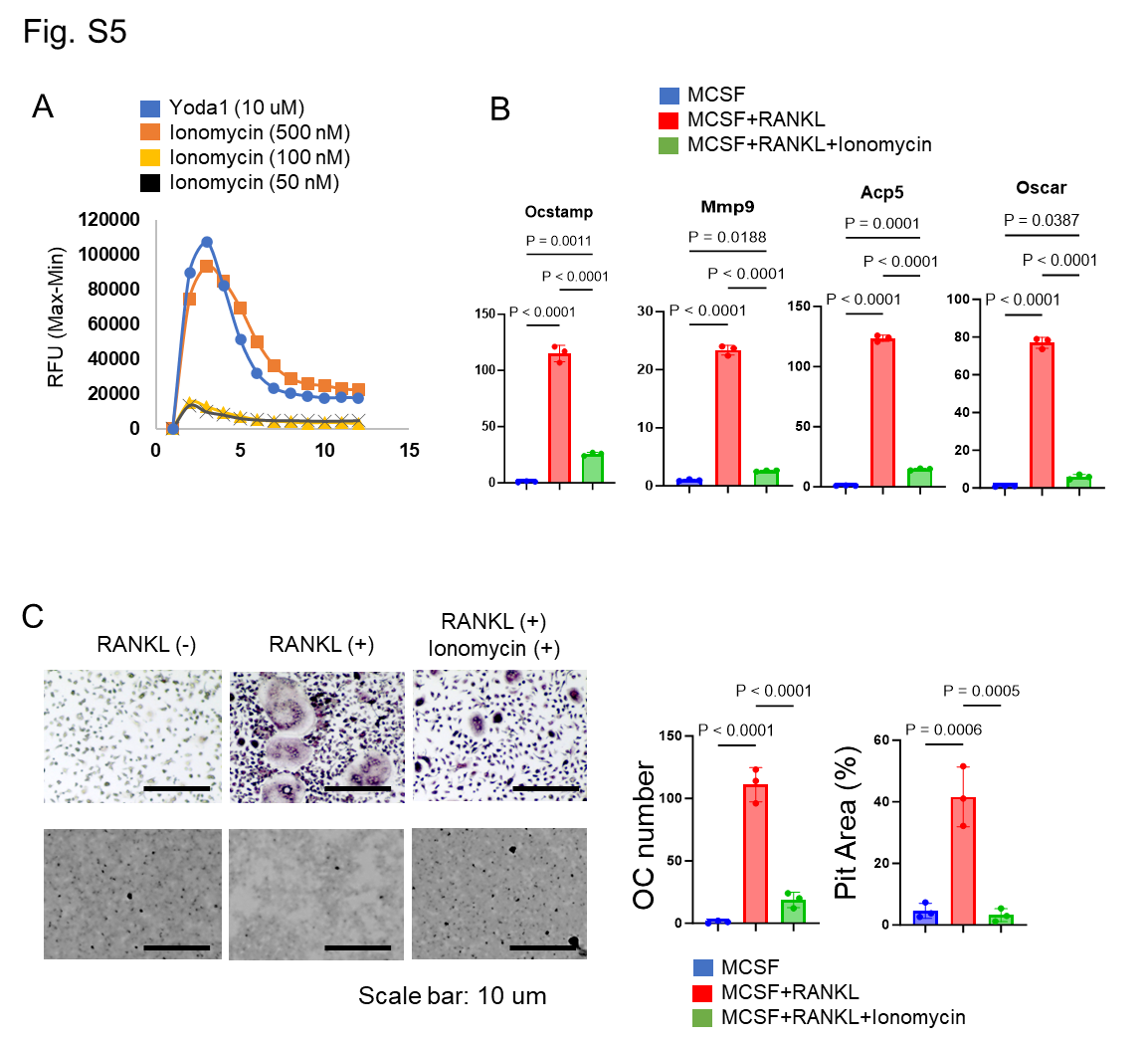

Figure S5. Ionomycin induces Ca^2+^ influx but inhibits OC-genesis.

500, 100 or 50 nM of Ionomycin were treated pre-OC to measure Ca^2+^ influx. (A) Ionomycin (100 nM) was applied to RANKL-mediated OCs to investigate the level of OC-related gene expression, such as Ocstamp, Mmp9, Acp5 as well as Oscar. (B) TRAP staining was conducted to evaluate Ionomycin (100 nM)-mediated OC-genesis. (C) Data represent the mean ± SD of three independent experiments.

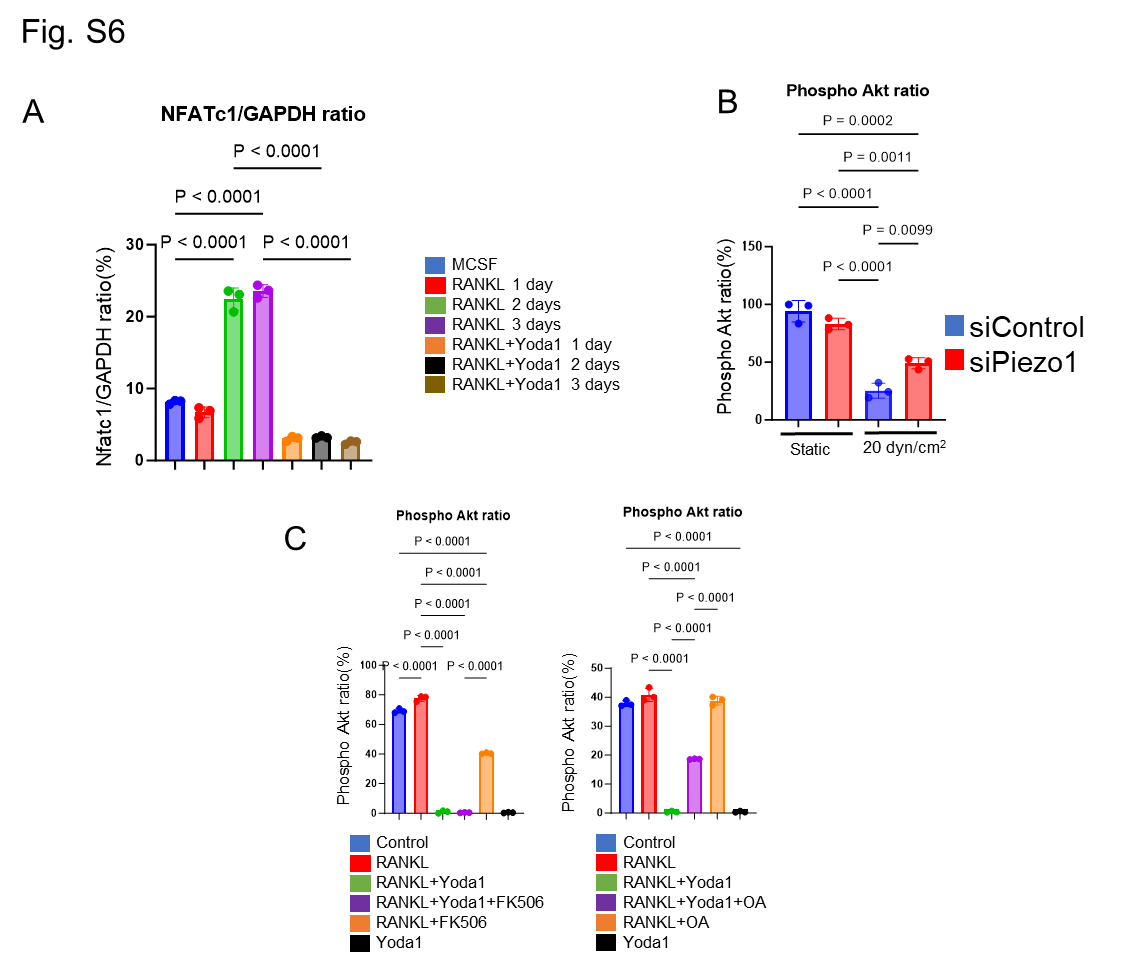

Figure S6. Densitometric analysis for western blotting.

Densitometric analysis of Western blotting in Fig. 3C, 5C and 6A was conducted using ImageJ software (Version 1.50). Data represent the mean ± SD of three independent experiments. Results were presented as the means ± SD.
